# The accidental ally: Nucleosomal barriers can accelerate cohesin mediated loop formation in chromatin

**DOI:** 10.1101/861161

**Authors:** Ajoy Maji, Ranjith Padinhateeri, Mithun K. Mitra

## Abstract

An important question in the context of the 3D organization of chromosomes is the mechanism of formation of large loops between distant base pairs. Recent experiments suggest that the formation of loops might be mediated by Loop Extrusion Factor proteins like cohesin. Experiments on cohesin have shown that cohesins walk diffusively on the DNA, and that nucleosomes act as obstacles to the diffusion, lowering the permeability and hence reducing the effective diffusion constant. An estimation of the times required to form the loops of typical sizes seen in Hi-C experiments using these low effective diffusion constants leads to times that are unphysically large. The puzzle then is the following, how does a cohesin molecule diffusing on the DNA backbone achieve speeds necessary to form the large loops seen in experiments? We propose a simple answer to this puzzle, and show that while at low densities, nucleosomes act as barriers to cohesin diffusion, beyond a certain concentration, they can reduce loop formation times due to a subtle interplay between the nucleosome size and the mean linker length. This effect is further enhanced on considering stochastic binding kinetics of nucleosomes on the DNA backbone, and leads to predictions of lower loop formation times than might be expected from a naive obstacle picture of nucleosomes.

## Introduction

The principles behind the organization of chromatin into a three-dimensional folded structure inside the nucleus remains an important open question [1–4]. An ubiquitous structural motif, as observed through Hi-C [5–8] and other experiments [9–11] are the formation of large loops, ranging from kilobases to megabases. These loops play both a structural as well as functional roles, bringing together regions of the DNA that are widely spaced along the backbone [12–14]. In recent years, much work has been done in trying to understand the mechanism of formation of these large loops. There is now a significant body of experimental observations that implicate a class of proteins called the Structural maintenance of chromosome (SMC) protein complexes - such as cohesin and condensin in the formation and maintenance of these large chromosomal loops [15–23].

Structural maintenance of chromosome (SMC) protein complexes are known to play a major role in chromosome segregation in interphase and mitosis. Both cohesin and condensin consists of SMC subunits and share structural similarities. SMC subunits (SMC1, SMC3 in cohesin and SMC2, SMC4 in condensin) fold back on themselves to form approximately a 50nm long arm. These two arms are then connected at one end by a hinge domain and other two ends which have ATPase activity are connected by a kleisin subunit (RAD21 in cohesin and condensin-associated protein H2 [CAPH2] in condensin)to form a ring like structure of the whole complex [24–30]. This ring like structure has been hypothesized to form a topological association with the DNA backbone [31, 32]. In particular, experiments on cohesin have shown that such topological association with the DNA can lead to very long residence times of cohesin [33]. The SMC ring can then embrace two chromosome strands, either within a single cohesin ring (embrace model), or within two cohesin rings mediated by an external protein (handcuff model) [30]. These two chromatin strands, bound topologically to the cohesin ring, can then extrude loops of chromosome, with the loop formation process ending at CTCF markers on the chromosome [30, 34–37].

There have been previous attempts to model this loop formation process through the action of SMC proteins. A common feature of these models is that the motion of these SMC proteins on the DNA backbone was assumed to be active, driven by the consumption of ATP [38–42], motivated by the presence of ATPase activity in the SMC proteins, and the fast timescales for the formation of these large loops. Such active SMC proteins have been theoretically shown to compact the chromosome effectively with stable loops formed by stacks of SMC proteins at the base of the loops [39]. The role of CTCF proteins in stopping the loop extrusion process has also been modeled and has successfully reproduced the occurrence of Topologically Associated Domains (TADs) in the simulated contact maps [37].

Recent experiments have however called into question this picture of active loop extrusion by cohesin. *In-vitro* experiments on DNA curtains have elucidated the nature of the motion of the cohesin protein on the DNA backbone. These experiments show that while the loading of the cohesin molecule on to the DNA is assisted by the ATP activity, the motion of the cohesin protein itself on the DNA strand is a purely diffusive process, and does not depend on ATP [33]. Analysis of the trajectory of cohesin on the DNA yields a time-exponent of 0.97, in excellent agreement with diffusive motion [33]. The measured diffusion coefficient of cohesin on bare DNA curtains is found to be *D* ≃ 1*µm*^2^/*s* at physiological salt concentrations of *c*_*KCl*_ ~ 100*mM* [33]. Further, the size of the cohesin ring implies that obstacles in the path of this diffusive trajectory can slow down the motion of cohesin. In particular, nucleosomes were found to act as obstacles to the diffusion of cohesin, and lowered the effective diffusion coefficient [33]. *In-vitro*experiments with a dense array of static nucleosomes have observed that the cohesin becomes almost static, and the estimated diffusive loop formation speeds at these high nucleosome densities was 7kb per hour [33], entirely too slow for the formation of the large loops that are seen in Hi-C experiments [10, 43–45]. In addition, While these certain external active proteins such as FtSz can drive cohesin actively along the backbone, the lifetime of these proteins are very small, unlike the topologically bound cohesin, and hence they cannot lead to persistent active motion of cohesin [33].

These experimental observations posit an interesting puzzle. Cohesin motion along the DNA appears to be purely diffusive, and the estimated diffusion coefficient seems incompatible with the formation of large loops. We investigate whether we can recover the fast loop formation times observed in experiments within the framework of passive, diffusive motion of cohesin. We show that the finite size of the nucleosome obstacles introduces an additional length scale in the system, and an interplay of this with the linker length can lead to non-monotonic looping times with varying nucleosome density. We report a regime where addition of nucleosomes can speed up the looping process, and we estimate looping times which explains how large loops may be formed even by a passively diffusing cohesin.

## Model

We consider only the one-dimensional diffusion of cohesin on chromatin. The DNA backbone is modeled as a one dimensional lattice of length *L*. The two subunits of the cohesin-chromatin complex (either within the same ring or within different cohesin rings) are modeled as two random walkers (RWs) that perform diffusive motion on this 1D lattice. The two cohesin subunits initally bind at neighboring sites on the DNA, and then start to drift apart due to diffusion. The length of the DNA between the subunits corresponds to the instantaneous size of the loop extruded. The two subunits cannot occupy the same site, with a loop of size zero corresponding to the situation when the subunits occupy neighbouring sites. The two ends of the DNA lattice correspond to the terminal points of the loop, and can biologically correspond to CTCF motifs which are known to act as endpoints for loop formation [30, 34–37]. In the context of our model, this is represented by absorbing boundary conditions at *x* = 0 and *x* = *L*.

Nucleosomes are modeled as extended objects in one dimension that cover and occlude *d* = 150 sites on the DNA lattice. Motivated by the experimental characterizations of motion of cohesin on nucleosome-bound DNA, we consider nucleosomes as barriers that reduce the local hopping rate of the cohesin rings. For a cohesin subunit present at a bulk site (no nucleosome on either side), the discrete master equation can then be written as,

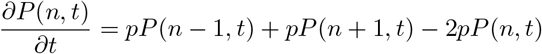

where, *P* (*n, t*) denotes the probability for the cohesin subunit to be at site *n* at time *t*. The hopping rate for cohesin in the bulk is denoted by *p*. At a site *r* which has a nucleosome to it’s right, the time evolution of the occupation probability can be written as,

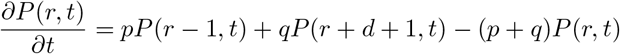

while for a site *l* that has a nucleosome to its left, we have,

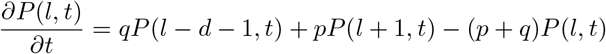

where *q* denotes the reduced barrier crossing rate of the cohesin ring in presence of a nucleosome (*q* ≪ *p*). Note that those hops which lead to both of the cohesin subunits occupying the same site are forbidden.

We are interested in determining the time taken by the cohesin molecule to form a loop of length *L*. The two rings absorb at the two boundaries at different times (*t*_*L*_ and *t*_*R*_) and the time taken to form the loop is defined as the maximum of these two times, *t*_*loop*_ = *max*{*t*_*L*_, *t*_*R*_}–this corresponds to the time when both the cohesin rings have reached the CTCF sites (at the lattice boundaries). This corresponds to the First Passage Time (FPT) for this stochastic problem of two random walkers in the presence of absorbing boundaries. We simulate the looping process on the discrete DNA lattice of length *L* using stochastic simulations (see *Methods*). Recent experiments on cohesin diffusion on DNA curtains has provided accurate estimates of the cohesin diffusion constant, *D*. We choose *D* = 1*µm*^2^/*s* corresponding to the experimental measurements at physiological salt concentrations [33]. This yields a hopping rate in the bulk of *p* = 9.1 × 10^6^/*s*. The reduced hopping rate when a cohesin ring has to pass through a nucleosomal barrier is given by *q* = 114*/s*, from experimental estimates of cohesin mobility [33].

## Results

### Nucleosomal barriers can accelerate cohesin looping

We first consider the case of static nucleosomes, i.e. when the nucleosomes occupy fixed random positions on the DNA lattice. The results for the looping time, *t*_*loop*_, and the effective diffusivity *D*_eff_ = *L*^2^/2*t*_*loop*_ as a function of the linker length (∆) are shown in Figs. 2 and 3 for a lattice of length *L* = 30*kbp*. Starting from the completely empty lattice (∆ = *L* = 30*kbp*), as we increase the number of nucleosomes (and hence decrease ∆), the loop formation time slowly increases (Fig. 2 black curve), and hence the effective diffusivity decreases (Fig. 3 black curve). This is expected since the nucleosomes act as extended barriers to the diffusion, and hence increasing the number of nucleosomes increases the time taken to form a loop. Contrary to naive expectations however, this increase in *t*_*loop*_ does not continue beyond a certain number of nucleosomes. Remarkably, beyond the point when the linker length becomes comparable to the nucleosome size itself, ∆ ≃ *d*, reducing the nucleosome spacing ∆ decreases the loop formation time, and hence increases the effective diffusivity. At the densest configuration of nucleosomes, the first passage time can reduce by two orders of magnitude from the slowest case at ∆ = *d*, and correspondingly, the diffusion coefficient can increase by two orders of magnitude.

**Figure 1.**
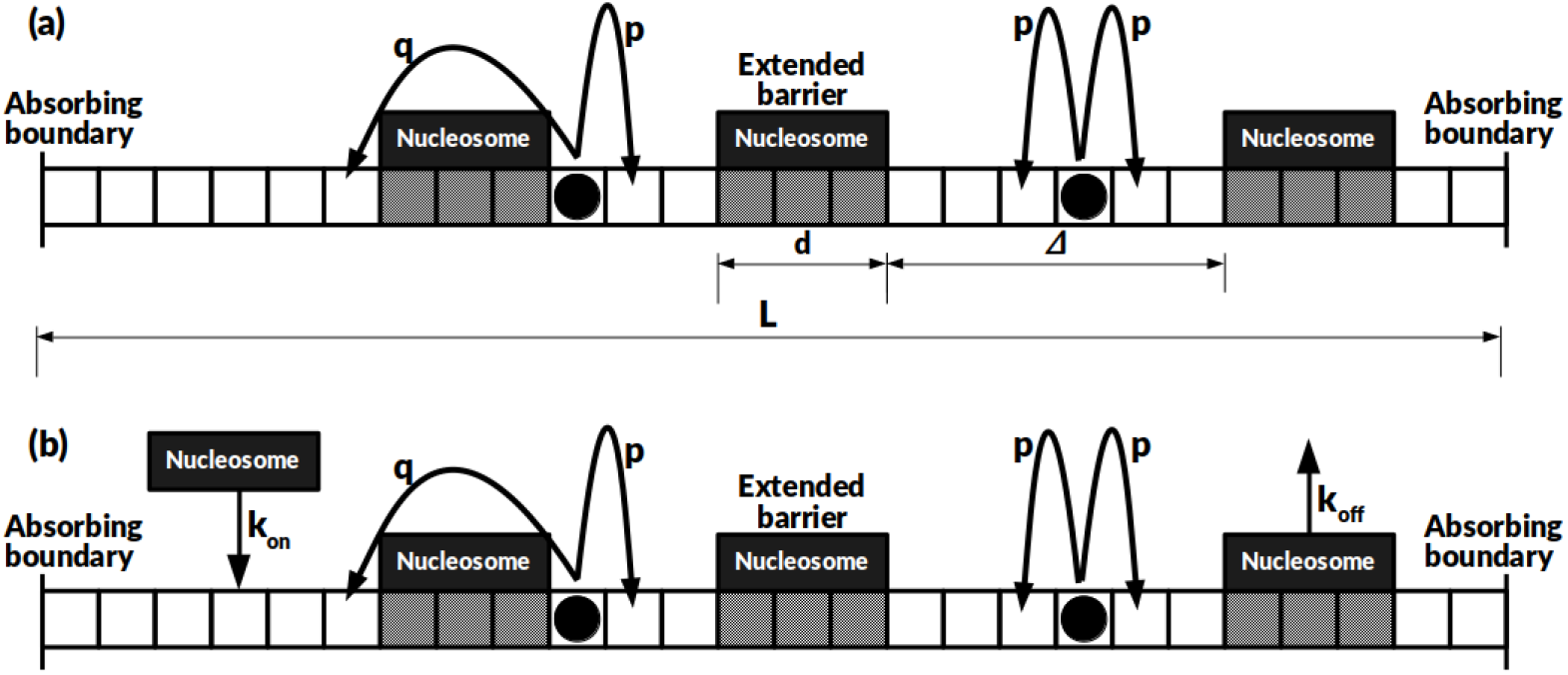
(a) Schematic of cohesin rings diffusing on DNA lattice. Nucleosomes form extended barriers covering *d* lattice sites. A cohesin ring traverses a nucleosomal barrier with a hopping rate *q* as compared to the bare lattice hopping rate *p*, with *q* ≪ *p*; (b) For the case of dynamic nucleosomes, nucleosomes can bind to the DNA lattice with a rate *k*_on_ and unbind with rate *k*_off_.

**Figure 2.**
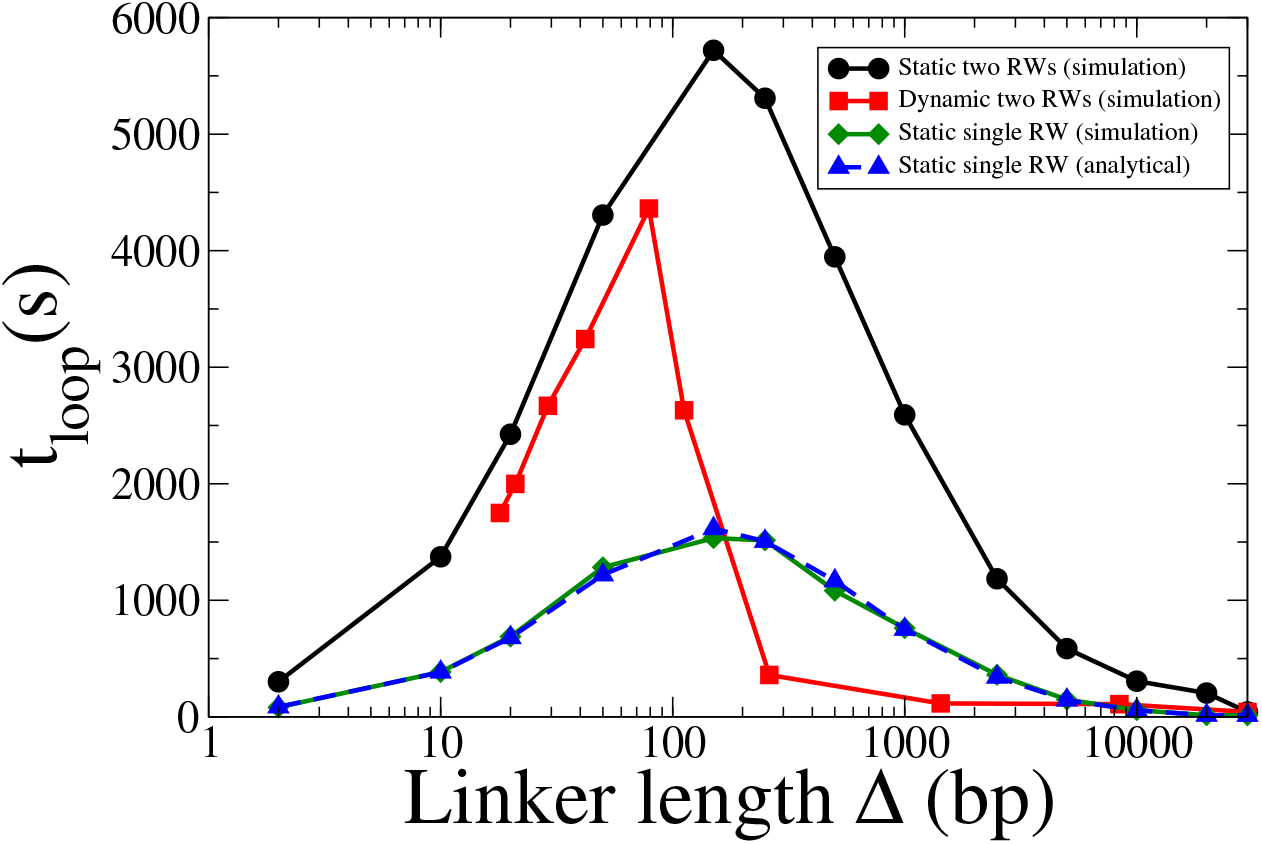
The loop formation time as a function of linker length for two cohesin subunits on a lattice with *L* = 30*kbp*. The black curve (circles) corresponds to results for two cohesin subunits in the presence of static nucleosomes. The blue (upper triangle) and green (diamond) curves correspond to the analytical and simulation results for a *single* subunit diffusing in the presence of static nucleosomes. Finally, the red curve (squares) shows the result for two cohesin subunits in the presence of dynamic nucleosomes. All cases show a non-monotonic dependence of the looping time on mean linker length, with a regime where looping times decreases with increasing nucleosome density.

**Figure 3.**
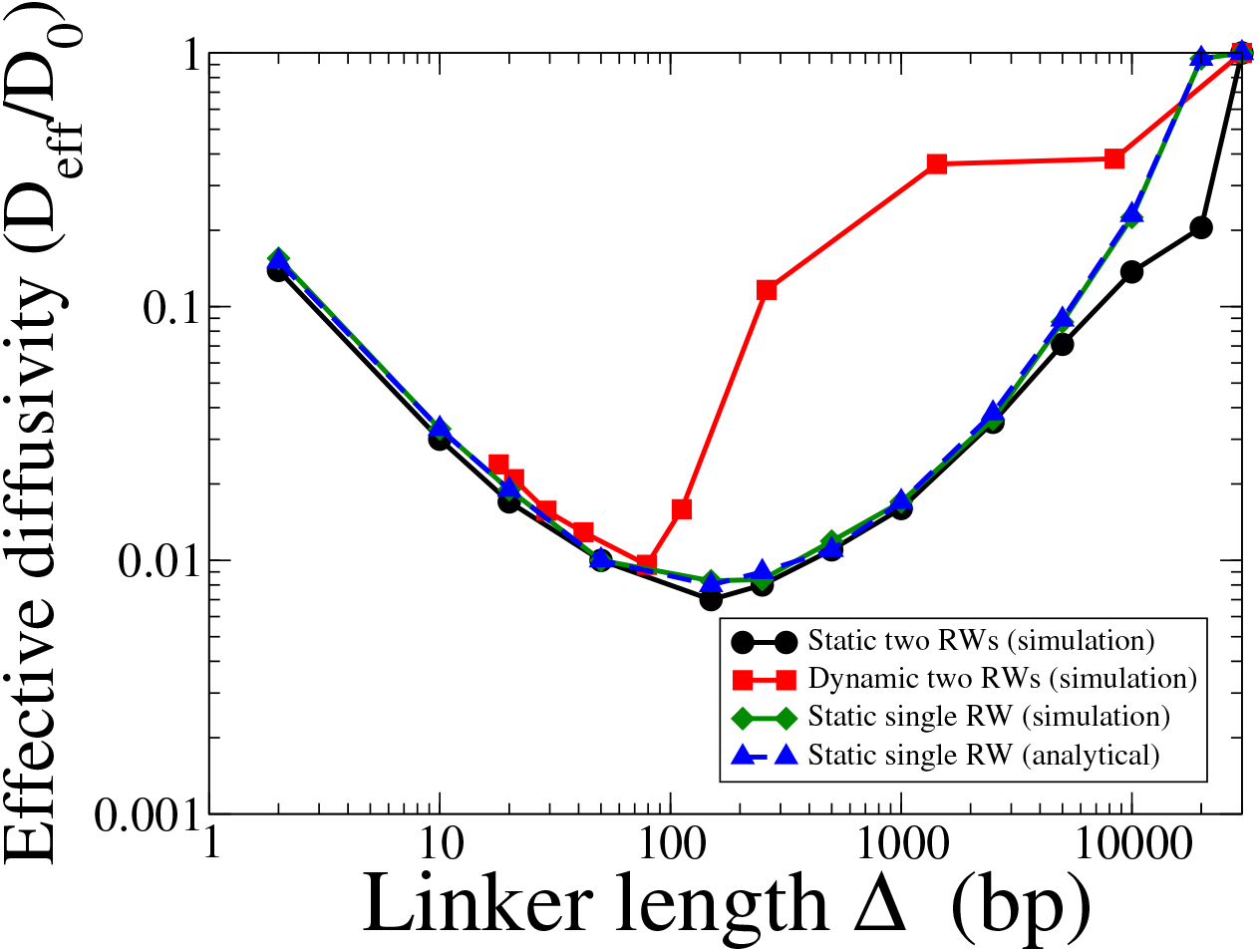
The normalised effective diffusivity, *D*_eff_/*D*_0_ as a function of linker length on a lattice with *L* = 30*kbp*. *D*_0_ represents the diffusivity on an empty DNA lattice (no nucleosomes). The different curves correspond to the same cases as described in Fig. 2. The non-monotonicity of the looping times is also reflected in the effective diffusivity, with a region where the diffusivity increases with increasing nucleosome number.

In order to obtain an theoretical understanding of the non-monotonicity of the loop formation times, and hence effective diffusivities, we solve the looping problem for a single cohesin subunit (RW). The mean looping time or the mean first passage time (MFPT) for a single diffusing walker starting from a bulk site *k* in a lattice of length *L*, *T*_*k,L*_ can be written as a recursion relation,

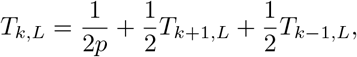

while, for a site that has a nucleosome to the right,

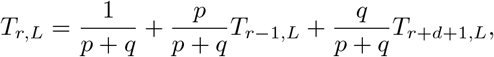

and for a site *l* that has a nucleosome to the left,

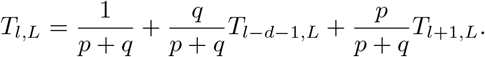

These recursion relations for the first passage times can be solved numerically subject to the absorbing boundary conditions,

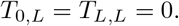

The comparison of the theoretical results with our simulations for a single RW is shown in Fig. 2 (dashed blue curve and solid green curve respectively). As can be seen from the figure, a single RW displays a similar non-monotonocity as in the case of two RWs, and this behaviour is captured by the analytical results for the looping time. The corresponding plots for the effective diffusion coefficient are shown in Fig. 3 (dashed blue curve and solid green curve). The non-monotonicity of the looping time and hence the effective diffusion coefficient is thus an integral feature of this random walk process in one dimension in the presence of extended barriers.

Physically, this can be understood as follows - the extended barriers pose an effective energy barrier that the cohesin subunit must overcome. Competing with this energy cost, there is also the entropic cost that is associated with the hopping of cohesin on the linker region between two nucleosomes. The effective free energy barrier is highest when the length of the nucleosome is comparable to the mean length of the linker DNA, leading to large escape times in this region. For linker lengths smaller than this critical value, the attempt rate for barrier crossing increases as the linker region shrinks, leads to faster barrier crossings and hence smaller looping times. This was verified explicitly by changing the nucleosome size in our simulations, and the largest loop formation time was always obtained when the mean linker length was equal to the assumed nucleosome size.

In addition to the mean loop formation time, we also calculate distributions of looping times. The distributions are shown in (Fig. 4) for three different values of the linker length. The looping time distribution is unimodal, with a peak at a finite looping time, and falls off exponentially as *t* → ∞. The non-monotonic nature of the mean looping times is also reflected in the full distribution, with the peak of the distribution being shifted to the right when the linker length becomes comparable to the size of the nucleosome, as can be seen for the case of ∆ = 150*bp* in Fig. 4(red curve). For linker lengths either smaller or larger than the nucleosome size, the distribution shifts to the left, commensurate with the observation of smaller loop formation times in these cases. This is a generic feature for this problem of 1D random walks with extended barriers in one-dimension, and continues to hold true for a single random walker.

**Figure 4.**
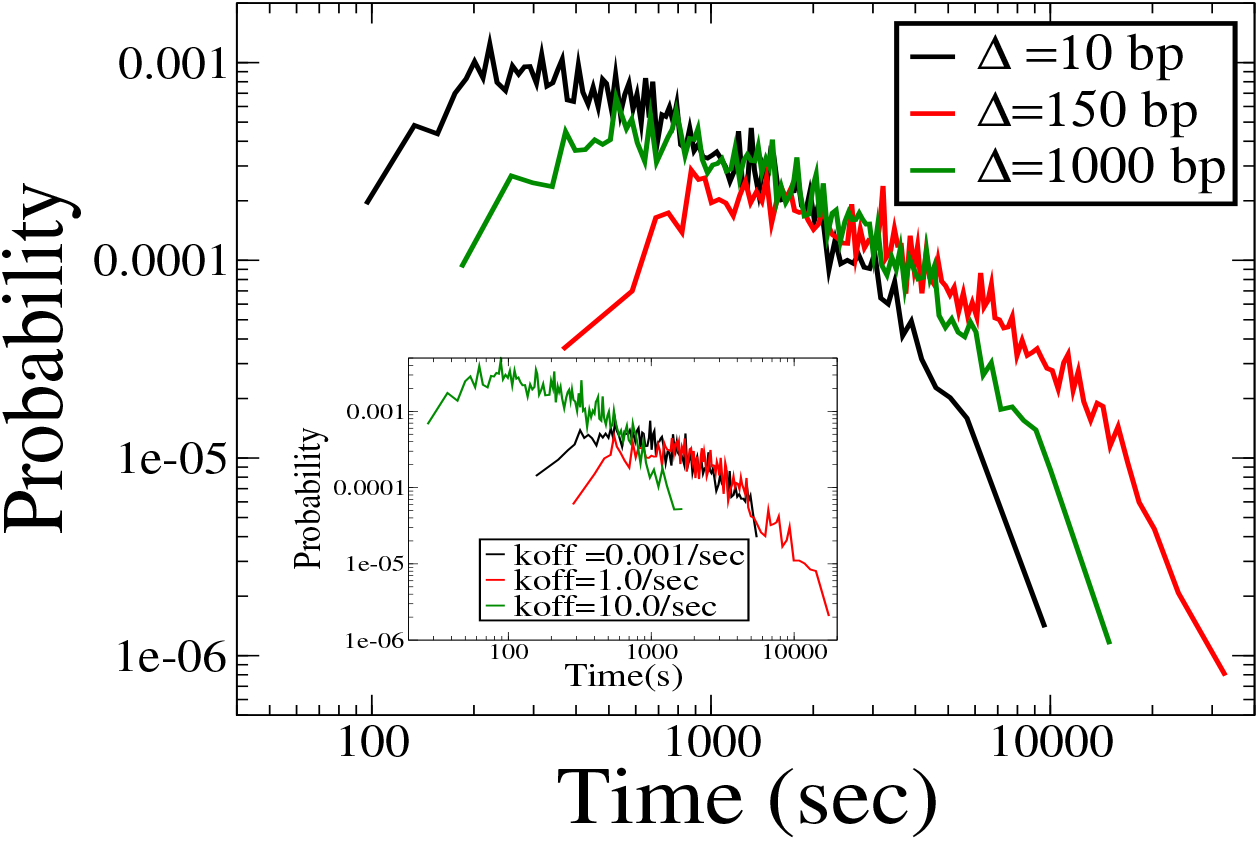
Distributions of loop formation times for three different values of inter-nucleosome spacing for the case of static nucleosomes for two RWs on a lattice of *L* = 30*kbp*. The distribution for ∆ = *d* is the broadest, consistent with the mean *t*_*loop*_ being the highest in this case. The inset shows the distributions for the case of dynamic nucleosomes for three different unbinding rates, and again the distribution for *k*_off_ = 1/*s* is the broadest.

### Nucleosome (un)binding kinetics can decrease loop formation times

We now turn to the case of dynamic nucleosomes, which can stochastically bind and unbind to the DNA lattice. Inside the nucleus, nucleosomes are dynamic and can regulate their binding and unbinding rates in response to gene activity. It hence becomes important to estimate the loop formation times by cohesin in the context of dynamic nucleosomes.

The loop formation times for the case of dynamic nucleosomes are consistently smaller than that obtained for static nucleosomes, and the first passage time can be reduced by as much as two orders of magnitude. For different values of *k*_off_, we obtain the mean linker length ∆(*k*_off_). Higher values of *k*_*off*_ correspond to lower nucleosome densities and hence larger linker lengths. We show the variation of the loop formation time with this mean linker length for dynamic nucleosomes in Fig. 2 (red curve). The non-monotonicity of the loop formation time persists, as in the case of static nucleosomes. However, the peak of the FPT curve is now shifted to smaller values of ∆, with the maximum loop formation time occuring around ∆ ≃ 50*bp*. The mean looping time drops sharply as the *k*_*off*_ (or equivalently, mean ∆) is increased, and approaches the empty lattice value for mean separations as small as ∆ ≈ 200*bp*.

This efficient speedup of the loop formation process is also apparent on considering the mean diffusivity, as shown in Fig. 3. The diffusivity drops only by two orders of magnitude even at the ∆ corresponding to the slowest loop formation times (∆ ≃ 50*bp*). At ∆ = 200*bp*, the effective diffusivity is *D*_*eff*_ ≃ 0.1*D*_0_. The distributions of looping times is also consistent with the non-monotonic nature of the mean looping. The inset of Fig. 4 shows the looping time distribution for three different values of *k*_off_. The distribution is broader for *k*_off_ = 1/*s* (red curve) than it is for unbinding rates both smaller (*k*_off_ = 0.001/*s*, black curve) and larger (*k*_off_ = 10/*s*, green curve) than this value. This mechanism thus provides a route to correlate gene activity of a chromatin segment to its loop formation efficiency, via the variations in the mean separation between nucleosomes.

### Statistical positioning of nucleosomes can marginally accelerate looping

It is well known that the stochastic binding and unbinding of nucleosomes results in an oscillatory occupancy profile from the start of the Transcription Start Site (TSS) [46, 47]. We assume, consistent with several experimental studies, that the TSS are correlated to the CTCF markers which define the endpoints of the loop [30, 34–37]. This oscillatory profile can be interpreted as an effective potential landscape in which the cohesin random walker executes its diffusive dynamics.

In order to determine whether the dynamic nature of the nucleosomes itself or the effective potential landscape imposed by the spatial variations of nucleosomal occupancy profiles ahead of a TSS is responsible for the predicted increase in the effective diffusivity, we investigate the variations of loop formation times on a lattice with periodic boundary conditions. The characteristic oscillatory profile of nucleosomal occupancy arises due to the finite boundaries at the ends of the lattice (CTCF markers), and hence is absent for the case of periodic lattice. This was explicitly verified for dynamic nucleosomes on a ring, where the nucleosomal occupancy shows a flat profile without any oscillations (see Fig. 5 inset).

**Figure 5.**
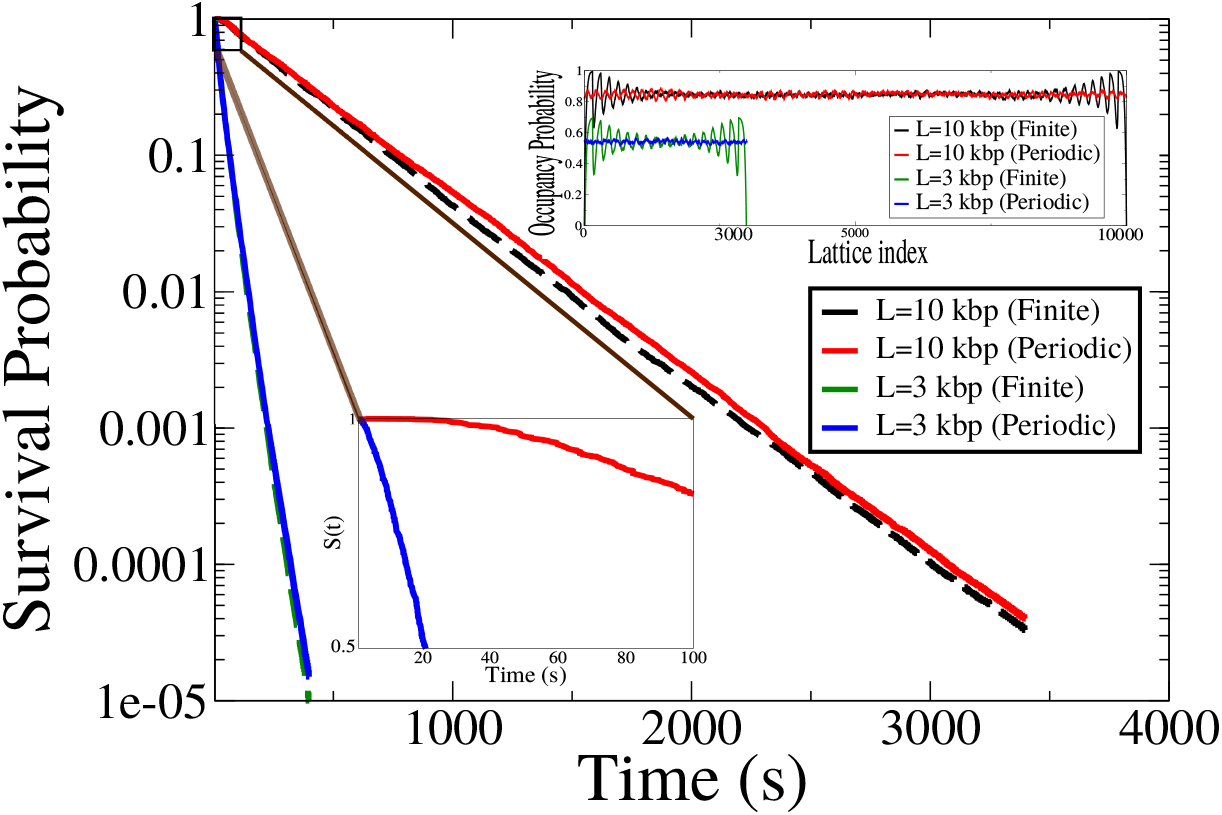
The effect of statistical positioning on mean looping times. Shown are the plots of the survival probabilities for a finite lattice (with statistical positioning, dashed lines) and a periodic lattice (no statistical positioning, solid lines), for two different lattice sizes, *L* = 3*kbp* and *L* = 10*kbp*. The unbinding rate is *k*_off_ = 0.01/*s*. The zoomed view illustrates the deviation of the survival probability from a single exponential for short looping times. The inset shows the nucleosomal occupancy probability as a function of the lattice site for the finite and periodic lattices. The plots for *L* = 3*kbp* have been vertically shifted so as to avoid overlap with the *L* = 10*kbp* plots. As expected, the periodic lattice shows no effect of statistical positioning.

We plot the survival probability *S*(*t*), defined as the probability that at least one of the two cohesin subunits survives till time *t*, as a function of time for a finite lattice and for a periodic lattice for two different lattice sizes (*L* = 3000bp and *L* = 10000bp) for a nucleosome unbinding rate of *k*_*off*_ = 0.01/*s*. We plot the survival probabilities using a ensemble cloning scheme (see *Methods* for details) in order to reliably access the tails of the distributions. As shown in Fig. 5, the difference in the survival probability distributions in the presence and absence of statistical positioning is relatively minor, showing that the nucleosome kinetics is primarily responsible for the small looping times for dynamic nucleosomes. However, the distribution for the case of periodic boundary conditions (when there is no effect of statistical positioning) consistently lies above the curve for the finite lattice. The mean looping times can be derived from the survival probability as, 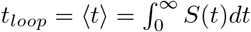. This gives 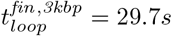 and 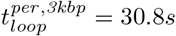, while for the 10*kbp* lattice, we obtain, 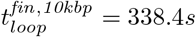 and 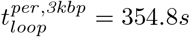, which also shows that the mean time for the periodic case is marginally higher than for the finite lattice. This implies that although the bulk of the speedup observed in the case of dynamic nucleosomes is due to the binding and unbinding of nucleosomes on the DNA backbone, statistical positioning can have a subtle effect in speeding up the formation of loops, and this can possibly become important for larger loop sizes.

### Loop formation time grows diffusively on loop length

We now turn to the question of how mean looping times scale with the size of the loop (lattice size), and the related question of how to characterize mean looping speeds. Previous analysis suggests that cohesin can spread diffusively on DNA over distances of 7*kb* in one hour [33].

Although our work suggests that looping time scales non-monotonically with the mean inter-nucleosome spacing (or equivalently, with *k*_*off*_ for dynamic nucleosomes), for a fixed value of ∆ (or *k*_*off*_), the mean loop formation time scales with the lattice size as *L*^2^, as expected from diffusive transport. This is shown on Fig. 6 for different values of ∆ for static nucleosomes, and for *k*_*off*_ = 100/*s* for dynamic nucleosomes.

**Figure 6.**
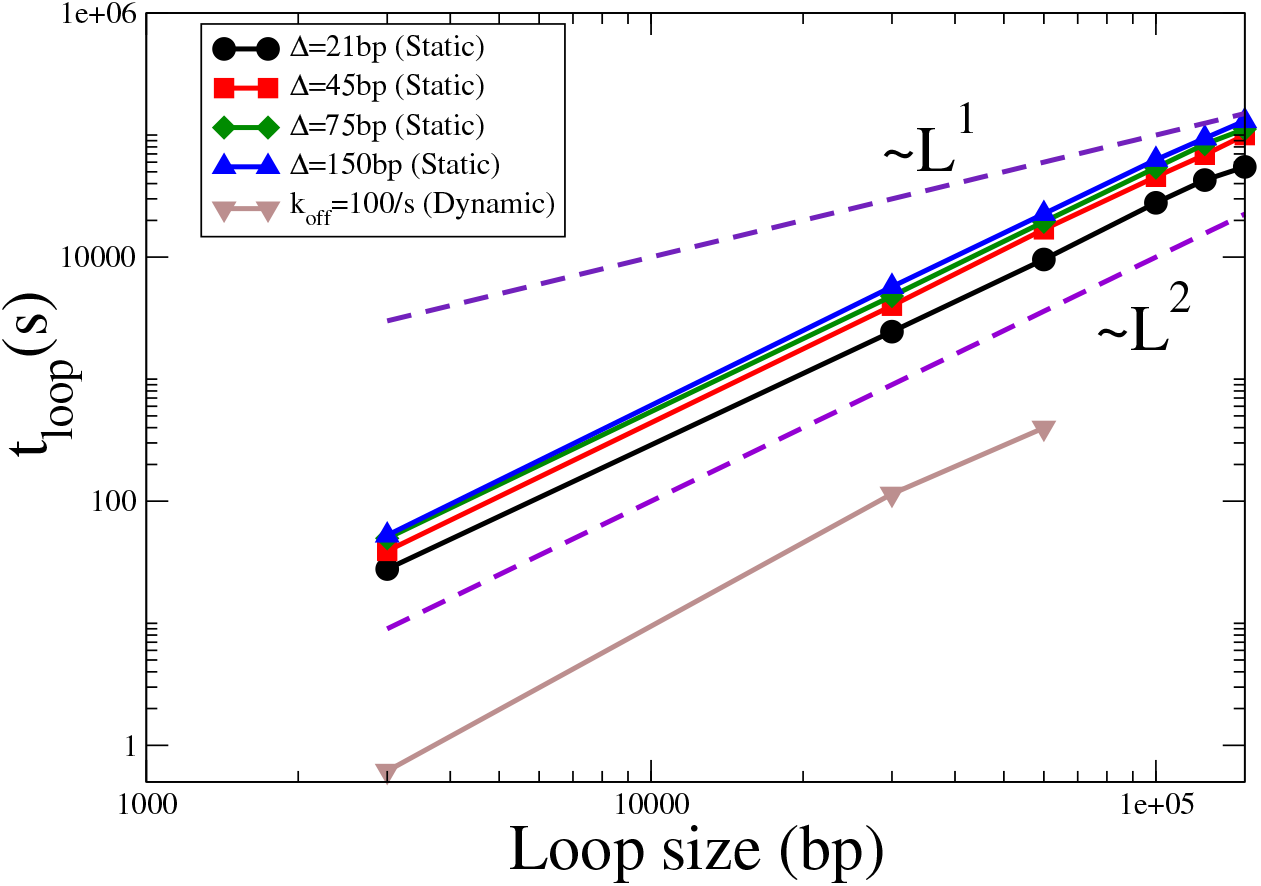
Scaling of the loop formation time with loop size for different inter-nucleosomal spacings, for static and dynamic nucleosomes. The *t* ∝ *L* and *t* ∝ *L*^2^ lines (dashed lines) are shown as guides to the eye. In all cases, the loop formation time grows diffusively with the size of the loop, consistent with the underlying diffusive dynamics.

Based on this analysis, we can now estimate looping speeds as predicted by this barrier-mediated diffusion model. For the case of static nucleosomes, at a mean nucleosome spacing of ∆ = 20*bp*, our analysis predicts that cohesin would form a 30*kbp* loop in a mean time of 2447*s*, corresponding to an effective looping speed of around 45*kbp* in one hour. For loop sizes of *L* = 100*kbp*, we obtain an effective looping speed of 13*kbp* in one hour. For dynamic nucleosomes, at *k*_*off*_ = 10/*s*, for a loop size of *L* = 30*kbp*, we obtain a mean looping speed of 300*kbp* per hour, a greater then 40−fold speedup as compared to the 7*kbp* per hour speeds calculated earlier.

## Discussion

We show that contrary to naive expectations of reduced diffusivities when cohesin faces a nucleosomal barrier, the true picture is far more nuanced. The macroscopic manifestation of *extended* microscopic barriers depends on the inter-barrier separation, the mean linker length. While the mean loop formation time initially increases with decreasing linker length, below a certain critical linker length (∆ ≲ *d*), this trend reverses, with the mean time now decreasing with decreasing linker lengths. At physiological nucleosome separations, this can lead to speed-ups of around two orders of magnitude compared to the diffusive speed estimated in previous studies. This counter-intuitive result is due to the extended nature of the nucleosomal barriers, which introduces an additional length scale in the system that competes with the inter-nucleosome spacing to give rise to this non-monotonic behaviour of the mean looping times.

This non-monotonic dependence of the looping time with linker length forms a testable prediction of our model. At extremely high densities of nucleosomes, our analysis predicts effective diffusivities that are comparable to the empty lattice diffusivity. An important consistency check can be made by comparing with experimental observations of almost stationary cohesin at very high nucleosome densities. These observations were made for a DNA strand of 48.5*kbp* containing 10-50 nucleosomes [33]. These correspond to mean linker lengths between ~ 800 − 5000*bp*, in which regime our model predicts extreme slowdown of cohesin diffusion, consistent with observations. In order to verify the non-monotonic nature, we would need to observe an even higher density nucleosomal array, with ≳ 200 nucleosomes on a 48.5*kbp* DNA strand.

Further, we illustrate how binding and unbinding of nucleosomes from the DNA backbone, can speed up diffusion of cohesin by upto two orders of magnitude compared to the static case. There is widespread experimental evidence that cells can tune the binding-unbinding kinetics of nucleosomes in response to different signals, and in general, active genes are characterised by more dynamic nucleosomes [48–51]. We show that this offers the cells a route to controlling the speed of loop formation by varying the nucleosome kinetics and hence linker length. This dependence of the looping time on the nucleosome kinetics also offers a tantalizing possibility of introducing directionality in the cohesin motion through an underlying asymmetry in the nucleosome positioning. Recent studies have opened the possibility of asymmetric nucleosome distributions near CTCF sites [52], and such an asymmetry can bias the underlying landscape in which cohesin performs its diffusive motion, leading to an effective drift term. While we have not explicitly investigated the effect of such an asymmetry in the current work, our results in Fig. 2 and Fig. 3 suggest such an asymmetry can further decrease looping times in real biological scenarios.

In addition to the effect of this extended barrier outlined in our work, several other factors may play a role to decrease looping times. Experiments have found while the motion of cohesin on the DNA backbone is itself diffusive, cohesins can be transiently pushed along the DNA by other active DNA motor proteins, such as FtSz [33]. Although this active push is short lived, as the FtSz protein changes direction, it can result in additional speed up of the loop formation process. Additionally, a recent theoretical proposal argues for a novel collective ratchet effect driven by a 1*D* osmotic pressure that favours the extrusion of larger loops [53, 54]. The mechanism underlined in this work stands apart from these other proposals in that it highlights the non-trivial role of the extended nucleosomal barriers. Other factors such as motor activity or collective effects would then serve to provide additional speed-ups to the estimates calculated in the present work.

Our works opens up a tantalizing possibility for the case of binding site search on DNA by a generic DNA-binding protein (DBP). The question of how DBPs search for target sites has a long history, with the leading hypothesis being that of facilitated diffusion, a combination of 3*D* and 1*D* diffusion [55–60]. During the phase of one-dimensional diffusion of the DBP on the DNA backbone, the proteins encounter nucleosomes. Although the specific topological association of cohesin on DNA may not be applicable to general proteins, one can imagine a DBP unbinding from the backbone and then re-attaching past the nucleosome site, which would then correspond to an effective reduced barrier crossing rate in the context of our model. The same physical phenomenon as outlined in this paper would then also be applicable, with the finite length of the nucleosomal barrier leading to effective speed-ups for certain linker lengths. This can result in faster 1D-search leading to lower effective search times.

In summary, our work highlights the non-trivial role of extended nucleosome barriers on the diffusion of cohesin on DNA. The extended barriers introduce an additional lengthscale which can reduce looping times beyond certain critical nucleosome densities. This non-trivial acceleration of DNA looping may serve to explain how cohesin forms large chromosomal loops even though it moves passively on the DNA backbone.

## Materials and Methods

### Static nucleosomes

In case of static nucleosomes, nucleosomes are placed on the DNA lattice maintaining a certain constant linker length (∆) between two consecutive nucleosomes. We verified that our results do not change if the nucleosomes are positioned randomly keeping the mean linker length to be the same. The two cohesin subunits, modeled as two RWs are initialized at two consecutive lattice sites near the middle of the lattice. At each timestep, we first choose one of the two subunits randomly, and update its position in accordance with the hopping rates p (if the subunit is not adjacent to a nucleosome) or q (if the subunit occupies a site adjacent to a nucleosome). We then repeat this for the other cohesin subunit. The two subunits are not allowed to occupy the same lattice site. We record the times *t*_*L*_ and *t*_*R*_ when the left and right RWs get absorbed at the boundaries. The looping time *t*_*loop*_ is the maximum of these two times. The simulation is repeated for ~ 1000 ensembles in order to obtain the mean looping time.

### Dynamic nucleosomes

In the case of dynamic nucleosomes, nucleosomes bind and unbind to the DNA lattice stochastically. We choose a fixed binding rate (*k*_*on*_) = 12/s [50, 61, 62] while the unbinding rate (*k*_*off*_) is varied in order to achieve different mean linker lengths. We first allow the system to reach a steady state nucleosomal occupancy in the absence of cohesin. Once the system reaches steady state, we position the cohesin subunits near the midpoint of the lattice. At each timestep, we choose the N+2 entities (N number of nucleosomes and two RWs) in random order. If a bound nucleosome is picked, it can unbind from the DNA with a rate *k*_*off*_; if an unbound nucleosome is picked, it can bind to the DNA with a rate *k*_*on*_. If either of the cohesin subunits are picked, they hop to an adjacent empty lattice site with a rate *p* or hop across a nucleosome with a rate *q*. The system evolves until both the cohesin subunits are absorbed at the two lattice boundaries. We again note the looping time *t*_*loop*_ and the mean looping time is obtained after averaging over *∼* 1000 such ensembles, as before.

### Ensemble cloning

In order to access the tails of the first passage (and survival probability) distributions, we use an ensemble cloning scheme, which can access these regions to a high degree of accuracy [63, 64]. As shown in Fig. 7, we start with 1000 distinct initial configurations at *t* = 0 and then allow all the 1000 systems to evolve for a time *T*. After this time *T*, in some systems both the RWs reach the end points of the lattice and are absorbed yielding a looping time *t*_*loop*_. In the remaining systems at least one of the RWs survives. We then clone these surviving systems to bring back the total number of systems to 1000 and again evolve for a time *T*. We repeat this procedure until we reach a desired accuracy for the survival probability.

**Figure 7.**
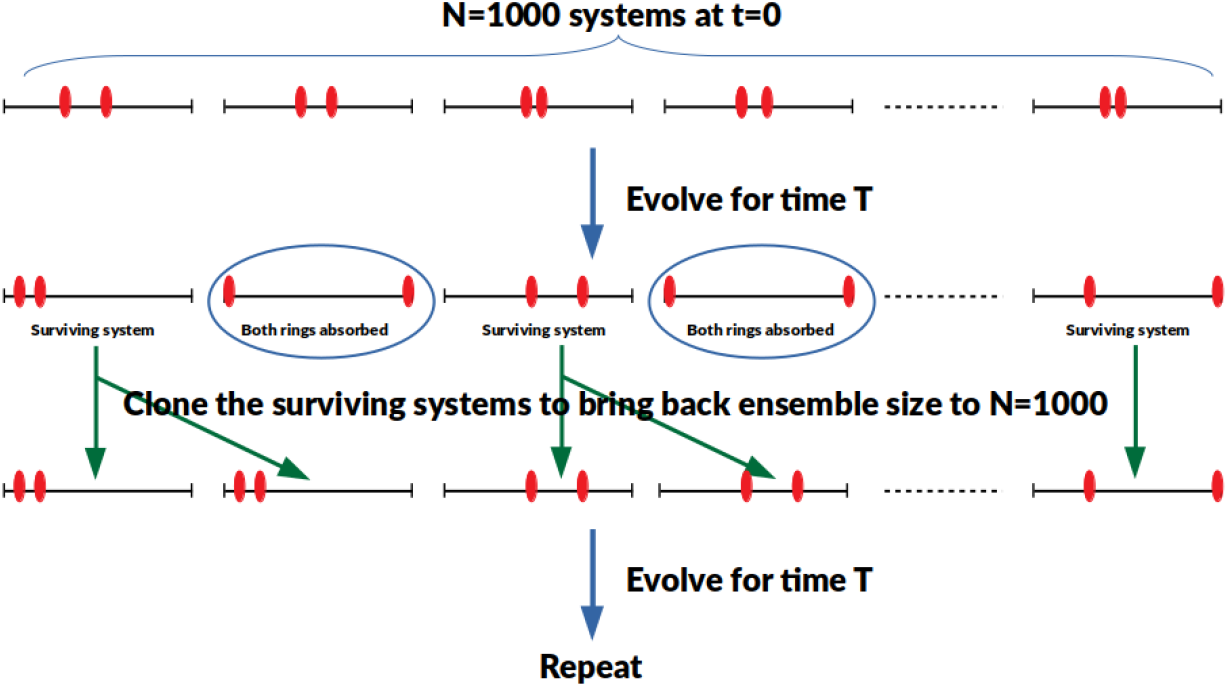
Schematic of the ensemble cloning scheme used to estimate survival probabilities.

At *t* = 0, the survival probability is *S*(0) = 1. Let, *M* (*T*) denote the number of systems which survive after evolving for time *T*. Then the survival probability at time *T* is, *S*(*T*) = *S*(0)(*M* (*T*)/1000). At this point, we clone these *M* (*T*) surviving systems to bring back the number of systems to 1000 and allow the systems to evolve for another interval of time *T*. Let, after this second iteration, the number of surviving systems be *M* (2*T*). Hence, the survival probability after this second iteration becomes, *S*(2*T*) = *S*(*T*) *×* (*M* (2*T*)*/*1000). The time *T* is chosen such that roughly half the number of systems survive after each iteration.

Thus the sruvival probability after *k* iterations is of the order of *∼* 2^*−k*^, and hence this procedure allows one to access the tails of the survival probability distribution.

## Acknowledgments

MKM acknowledges funding support from the Ramanujan Fellowship (13DST052), DST and IIT Bombay (14IRCCSG009).

## Author contributions

RP and MKM designed and supervised research. AM performed research. AM and MKM analyzed data. MKM supervised analysis. AM, RP and MKM wrote the paper.

